# Characterization of Human Dosage-Sensitive Transcription Factor Genes

**DOI:** 10.1101/528554

**Authors:** Zhihua Ni, Xiao-Yu Zhou, Sidra Aslam, Deng-Ke Niu

## Abstract

Copy number changes in protein-coding genes are detrimental if the consequent changes in protein concentrations disrupt essential cellular functions. The dosage sensitivity of transcription factor (TF) genes is particularly interesting because their products are essential in regulating the expression of genetic information. From four recently curated datasets of dosage-sensitive genes (genes with conserved copy numbers across mammals, ohnologs, and two datasets of haploinsufficient genes), we compiled a dataset of the most reliable dosage-sensitive (MRDS) genes and a dataset of the most reliable dosage-insensitive (MRDIS) genes. The MRDS genes were those present in all four datasets, while the MRDIS genes were those absent from any one of the four datasets and with pLI values < 0.5 in both of the haploinsufficient gene datasets. Enrichment analysis of TF genes among the MRDS and MRDIS gene datasets showed that TF genes are more likely to be dosage-sensitive than other genes in the human genome. The nuclear receptor family was the most enriched TF family among the dosage-sensitive genes. TF families with very few members were also deemed more likely to be dosage-sensitive than TF families with more members. In addition, we found a certain number of dosage-insensitive TFs. The most typical were the KRAB domain-containing zinc-finger proteins (KZFPs). Gene Ontology (GO) enrichment analysis showed that the MRDS TFs were enriched for many more terms than the MRDIS TFs; however, the proteins interacting with these two groups of TFs did not show such sharp differences. Furthermore, we found that the MRDIS KZFPs were not significantly enriched for any GO terms, whereas their interacting proteins were significantly enriched for thousands of GO terms. Although KZFPs and some other MRDIS TFs likely have lower functional complexity than MRDS TFs, they were under-annotated in the GO database. We also examined whether the presence of a repressor domain and an activation domain underlies the mechanism of dosage sensitivity of TFs. Further characterizations revealed significant differences between MRDS TFs and MRDIS TFs in the lengths and nucleotide compositions of DNA-binding sites as well as in expression level, protein size, and selective force.

## 1 INTRODUCTION

Gene duplication and loss in evolution and gene copy number polymorphisms at the population level have been widely observed in both animals and plants (Innan and Kondrashov, 2010; Schrider and Hahn, 2010; Panchy et al., 2016). The copy number of a particular gene present in a genome is termed the gene dosage. A dosage change in a gene can happen within one locus through inactivation of one or two alleles or among two or more loci through gene duplication and loss of duplicates. Gene dosage sensitivity is the model of the deleterious effects resulting from genomic alteration of gene dosage (Rice and McLysaght, 2017a).

If the protein products of both alleles are required for a normal phenotype, heterozygous loss of function results in an abnormal phenotype. Haploinsufficiency, a measure of intolerance to such heterozygous loss-of-function variations, is a widely studied model of gene dosage sensitivity. Here, we briefly describe two sources of data on human gene haploinsufficiency used in this study. According to the sensitivity to loss-of-function variation, each human gene can be assigned to one of three natural categories: null (in which loss-of-function variation, regardless of whether it is heterozygous or homozygous, is completely tolerated by natural selection), recessive (in which heterozygous loss-of-function variation is tolerated but homozygous loss-of-function (HLOF) variation is not), and haploinsufficient (in which heterozygous loss-of-function variation is not tolerated). Lek et al. (2016) assumed that tolerant genes have the expected amount of truncating variation and took the empirical mean observed/expected rates of truncating variation for recessive disease genes and severe haploinsufficient genes to represent the average outcome of the homozygous and heterozygous intolerant scenarios, respectively; they then built a three-state model and designed a metric, the probability of being loss of function-intolerant (pLI). This metric can be used to evaluate the intolerance to heterozygous protein-truncating variation of each gene. With a cutoff of a pLI > 0.9, the researchers identified 3230 haploinsufficient genes by analyzing 60706 human exomes. In addition to previously identified haploinsufficient genes associated with diseases, their dataset included some genes that had not yet been assigned to any human diseases. The low incidence of these genes in human populations indicates that heterozygous loss of function confers some survival or reproductive disadvantage. More recently, Shihab et al. (2017) integrated genomic and evolutionary information from several large databases and predicted the existence of 7841 haploinsufficient genes in the human genome using a machine learning approach called HIPred. This dataset was comparably larger than that of Lek et al. mostly because Shihab et al. used a relaxed cutoff of a pLI > 0.5.

As the dosage sensitivity of a gene results from effects related to the ratio of its product to other cellular components, small-scale duplication events involving dosage-sensitive genes are deleterious. Therefore, the duplicability of a gene in evolution relative to all other genes in the same genome can also be used as a measure of dosage sensitivity (Rice and McLysaght, 2017a). Pairs of genes in the same genome originating from small-scale duplication are termed paralogs, while those originating from whole-genome duplications are termed ohnologs (Glover et al., 2016). If a gene has only ohnologs but no paralogs, its copy number changes during evolution are very likely under dosage constraints. Makino and McLysaght (2010) compiled a list of 7294 ohnologs in the human genome and confirmed that these ohnologs are in chromosomal regions with low copy number variations. Later, Rice and McLysaght (2017b) found that the copy numbers of genes in pathogenic copy number-variation regions of the human genome are more conserved across 13 mammalian genomes than those of genes in benign regions. Therefore, the 7014 genes they identified to have the most conserved copy numbers across the 13 mammalian genomes could be regarded as dosage-sensitive genes.

Transcription factors, in a narrow sense, are proteins that control the rate of transcription by binding to specific DNA sequences (Lambert et al., 2018). Mutations affecting the DNA-binding domains (DBDs) of TFs disrupt the accurate control of gene expression and cause diseases. It should be noted that some proteins that regulate gene expression but do not directly bind DNA are also called TFs. Previous studies have shown that some TFs work in a dose-dependent manner; heterozygous loss-of-function mutations of TFs are also associated with severe phenotypic disorders (Engelkamp and van Heyningen, 1996; Seidman and Seidman, 2002). In addition, in some cases, individuals with heterozygous deletions of particular TFs (e.g., THRB) do not exhibit abnormal phenotypes (Engelkamp and van Heyningen, 1996; Seidman and Seidman, 2002). A strong majority of genes in the human genome are dosage insensitive (Makino et al., 2013; Lek et al., 2016; Rice and McLysaght, 2017a; Rice and McLysaght, 2017b; Shihab et al., 2017). However, considering that the function of a TF is to ensure the expression of a target gene in the right cell at the right time and in the right amount in response to intracellular, intercellular or environmental signals, it seems odd for a TF to exist whose dose reduction or duplication does not cause any phenotypic changes. In 2002, Seidman and Seidman (2002) surveyed the molecular causes of human syndromes. Among 491 TF genes, only 27 were confirmed to exhibit phenotypic haploinsufficiency in the syndromic records available at that time. Members of several TF gene families appear more likely to be sensitive to copy number changes than members of other families. Are the previously identified dosage-insensitive TF genes (Engelkamp and van Heyningen, 1996; Seidman and Seidman, 2002) just exceptions, or is dosage sensitivity limited to certain unique families of TF genes? Recently, Lambert et al. (2018) curated a comprehensive list of human TFs by manually examining lists of putative TF from several sources, including previous manual curations, domain searches, the Gene Ontology (GO) database, and crystal and NMR structure data on proteins in complex with DNA taken from the Protein Data Bank. This most updated list includes 1639 TFs categorized into 65 families. Another study in the same year curated 1665 TFs, a very similar number (Hu et al., 2018). Taking advantage of an unprecedented wealth of data with regard to both the annotation of TF genes in the human genome and the identification of dosage-sensitive genes, we re-evaluated the dosage sensitivity of human TF genes. By comparing dosage-sensitive and dosage-insensitive TF genes, we characterized the mechanistic properties of the sensitive genes, including their DNA quaternary structures, DNA-binding sites, DBDs, behaviors in protein-protein interactions (PPIs), GO enrichments, expression patterns, selective forces (nonsynonymous difference [*d*_N_]/synonymous difference [*d*_S_] values) experienced in evolution, etc.

## 2 MATERIALS AND METHODS

A list of 1639 human TFs, their classifications into different families, their DNA-binding motifs, and their quaternary structures were obtained from Lambert et al. (2018). Data on the DBDs of human TFs also contributed by Lambert et al. (2018) were downloaded from http://humantfs.ccbr.utoronto.ca/download.php. Data on transactivation domains and transcriptional repressor domains were retrieved from the UniProt Knowledgebase (UniProtKB, UniProt release 2019_04) (The UniProt Consortium, 2019). Data on Krüppel-associated box (KRAB) domain-containing zinc-finger proteins (KZFPs) were obtained from Imbeault et al. (2017). Data on PPIs were obtained from the STRING database (Szklarczyk et al., 2018).

A list of 1317 HLOF-tolerant human genes was obtained from Saleheen et al. (2017). From Lek et al. (2016), we retrieved the probability of a gene to be tolerant to both heterozygous loss-of-function and HLOF mutations (pLI and pNull values) of all human genes. With a cutoff of > 0.9 for both metrics, 1226 HLOF-tolerant genes and 3230 haploinsufficient genes were obtained. A larger dataset of human haploinsufficient genes (7841) was obtained from Shihab et al. (2017). A list of 7294 ohnologs in the human genome was obtained from Makino and McLysaght (2010) and Makino et al. (2013), and a list of 7014 copy number-conserved genes was obtained from Rice and McLysaght (2017b). The orthologous relationships between human and mouse genes and their *d*_N_ and *d*_S_ values were retrieved from BioMart (Ensembl version GRCh37, http://grch37.ensembl.org/biomart/martview). Gene expression data were downloaded from the Human Protein Atlas (Version 18.1, https://www.proteinatlas.org/) (Uhlén et al., 2015), and the length of each protein-coding sequence was retrieved from Ensembl (version GRCh37, ftp://ftp.ensembl.org/pub/grch37/update/fasta/homo-sapiens/). Only transcripts with full open reading frames and multiples of three nucleotides were retained. For the multiple alternative splicing isoforms, we retained only the longest transcripts. In total, 22810 protein-coding genes have been annotated in version GRCh37.p13 of the human genome.

In enrichment analyses, chi-square tests (expected value > 5) and Fisher’s exact tests (expected value ≤ 5) were used to test whether a particular set of TF genes was significantly over- or underrepresented in a dataset of dosage-sensitive genes or HLOF-tolerant genes. As tens of chi-square or Fisher’s exact tests were performed per dataset, some of the obtained *p* values were likely to be less than 0.05 purely by chance. Therefore, the false discovery rate (FDR)-adjusted *p* values were computed using the Benjamini-Hochberg (BH) procedure.

Enrichments and comparisons of GO terms were performed using the R package clusterProfiler with its default settings (Yu et al., 2012). In this program, adjusted *p* values were also estimated to prevent a high FDR during multiple testing.

## 3 RESULTS

According to Lambert et al. (2018), 1639 annotated TF genes are in the human genome. Among them, 1570 TF genes were categorized into 64 families according to the DBDs they encoded. In addition, 69 TFs lacking recognizable DBDs were collectively categorized into a family termed “Unknown”. The largest family was the C2H2-ZF family, with 747 members, followed by the homeodomain (196 members), bHLH (108 members), bZIP (54 members), forkhead (49 members), nuclear receptor (46 members), HMG/Sox (30 members), ETS (27 members), T-box (17 members), AT-hook (16 members), Homeodomain+POU (16 members), Myb/SANT (14 members), THAP finger (12 members), CENPB (11 members), E2F (11 members), BED ZF (10 members), GATA (10 members), and Rel (10 members) families, among others. In addition, we were also interested in whether TF gene families with very few members have some uniqueness with regard to dosage sensitivity. For this reason, we defined four additional categories of small TF gene families, each with ≤ 5, ≤ 7, ≤ 9, or ≤ 11 members. All the TF members within each category were grouped together in statistical analyses like a gene family.

### 3.1 Enrichment of TF Genes among the Dosage-Sensitive Genes

Using a machine learning approach called HIPred, Shihab et al. (2017) predicted 7841 haploinsufficient genes in the human genome. Among them, 7824 genes have been annotated in version GRCh37.p13 of the human genome. By analyzing the variations across 60706 human exomes, Lek et al. (2016) identified 3230 haploinsufficient genes.

Ohnologs are duplicates of genes whose duplication can be tolerated only in whole-genome duplication events. Their duplicability is limited by their dosage relative to other genes of the same genome. Using an all-against-all blastp search of human, zebrafish (*Danio rerio*), green spotted puffer (*Tetraodon nigroviridis*), stickleback (*Gasterosteus aculeatus*), medaka (*Oryzias latipes*), Japanese puffer (*Takifugu rubripes*) and sea vase (*Ciona intestinalis*, an ascidian) protein sequences, Makino and McLysaght (2010) identified 7294 ohnologs in the human genome. Among them, 7074 genes have been annotated in the version GRCh37.p13 of the human genome.

In addition, Rice and McLysaght (2017b) identified 7014 human genes whose copy numbers are conserved across 13 mammalian genomes (*Bos taurus, Callithrix jacchus, Canis lupus familiaris, Equus caballus, Felis catus, Gorilla gorilla, Macaca mulatta, Mus musculus, Oryctolagus cuniculus, Ovis aries, Pan troglodytes, Rattus norvegicus* and *Sus scrofa*) and showed evidence that these copy number-conserved genes are dosage-sensitive.

We noticed that the datasets of dosage-sensitive genes obtained through different methods varied substantially (Figure 1). The dosage sensitivity of 853 genes was consistently supported by the four independent studies, so we regarded these genes as the most reliable dosage-sensitive (MRDS) genes (Supplementary Table 1). The MRDS gene dataset contained 122 TF genes representing 32 TF gene families. Statistical analysis showed that TF genes were significantly overrepresented in the MRDS gene dataset (chi-square = 20.1, df = 1, BH-adjusted *p* = 3 × 10^− 4^). This significant overrepresentation was observed in nine TF gene families, including the nuclear receptor, Grainyhead, bHLH, C2H2-ZF;Homeodomain, T-box, AP-2, RFX, Rel, and paired box families (BH-adjusted *p* < 0.05 for all cases, Table 1). The small TF gene families were also overrepresented in the MRDS gene dataset regardless of whether they were defined by ≤ 5, ≤ 7, ≤ 9, or ≤ 11 members (BH-adjusted *p* < 0.05 for all cases, Table 1).

**Table 1.** TF genes in the most reliable dosage-sensitive (MRDS) gene dataset.

| TF family | Total number in human genome | Observed number of MRDS TFs | Expected number in this dataset | BH-adjusted *p* |
| --- | --- | --- | --- | --- |
| Nuclear receptor | 46 | 20 | 1.72 | $1.5 \times 10^{-14}$ |
| Grainyhead | 6 | 4 | 0.22 | $6.0 \times 10^{-4}$ |
| C2H2-ZF;Homeodomain | 4 | 3 | 0.15 | 0.002 |
| T-box | 17 | 5 | 0.64 | 0.003 |
| AP-2 | 5 | 3 | 0.19 | 0.004 |
| bHLH | 108 | 12 | 4.04 | 0.005 |
| RFX | 8 | 3 | 0.30 | 0.016 |
| Rel | 10 | 3 | 0.37 | 0.030 |
| Paired box | 4 | 2 | 0.15 | 0.040 |
| All TFs | 1639 | 122 | 61.29 | $2.6 \times 10^{-4}$ |
| Small-family TFs |  |  |  |  |
| ≤ 5 members | 87 | 15 | 3.25 | 0.032 |
| ≤ 7 members | 126 | 23 | 4.71 | 0.004 |
| ≤ 9 members | 186 | 31 | 6.96 | 0.001 |
| ≤ 11 members | 238 | 37 | 8.90 | $5.9 \times 10^{-4}$ |

The chi-square test (expected value $> 5$ ) and Fisher's exact test (expected value $\leq 5$ ) were used to test the overrepresentation or underrepresentation of the TF genes in this dataset. The Benjamini-Hochberg (BH) procedure was used to compute the false discovery rate-adjusted $p$ values. The MRDS genes were defined as the common dosage-sensitive genes among the four datasets obtained by Shihab et al. (2017), Lek et al. (2016), Makino et al. (2013) and Rice and McLysaght (2017b).

**Figure 1.**
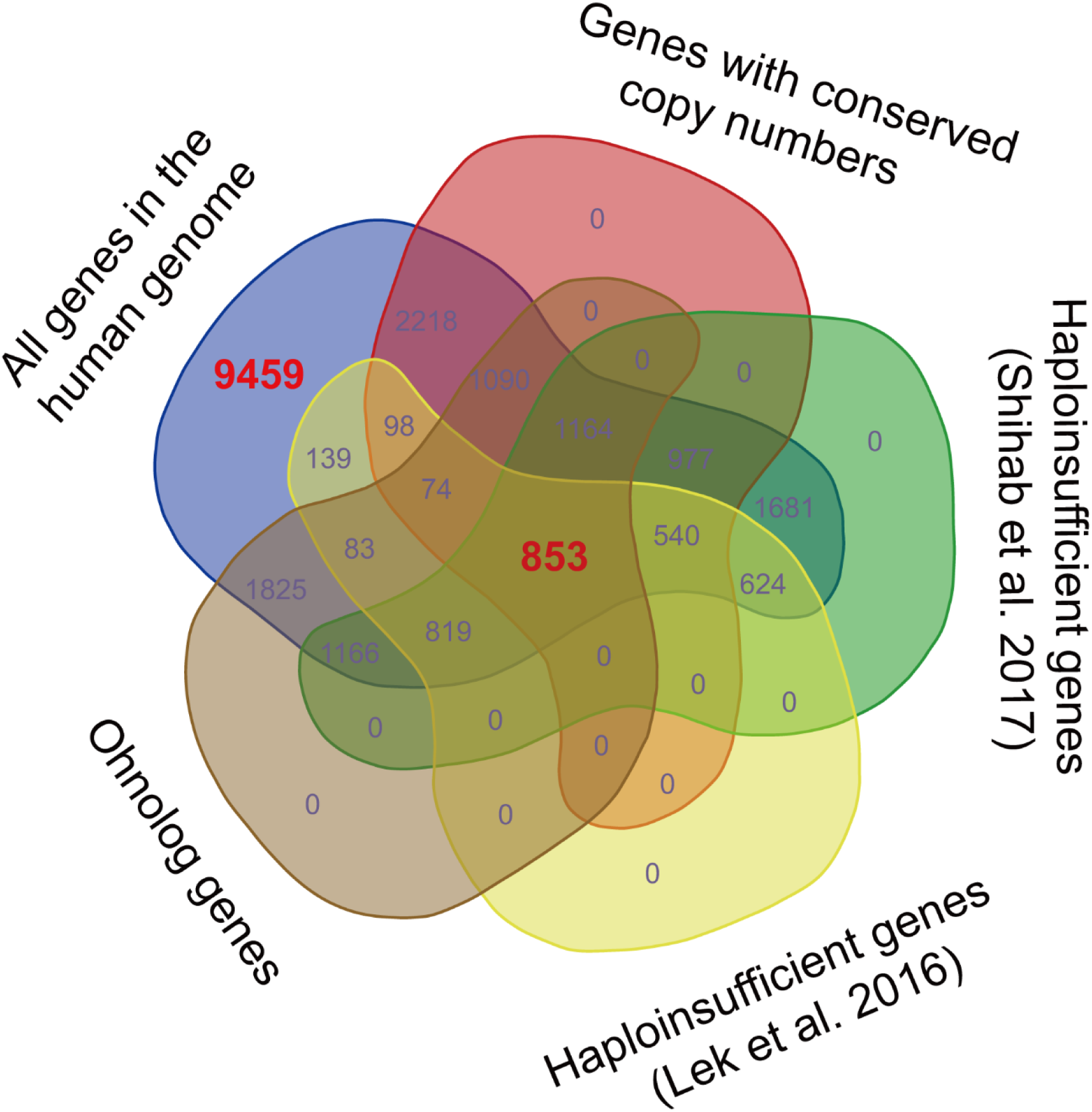
Venn diagram displaying the variations among dosage-sensitive gene datasets obtained through different methods. The 853 dosage-sensitive genes shared by the four datasets obtained by Shihab et al. (2017), Lek et al. (2016), Makino et al. (2013) and Rice and McLysaght (2017b) were regarded as the most reliable dosage-sensitive (MRDS) genes. To obtain a dataset of the most reliable dosage-insensitive (MRDIS) genes, the 9459 genes that were absent from any of the four datasets were further filtered by discarding the genes with a pLI value > 0.5 in either the dataset of Shihab et al. (2017) or the dataset of Lek et al. (2016). In total, 5579 MRDIS genes were obtained.

Furthermore, we observed 9459 genes that were not detected to be dosage-sensitive in any of the four studies on gene dosage sensitivity (Makino and McLysaght, 2010; Lek et al., 2016; Rice and McLysaght, 2017b; Shihab et al., 2017). To obtain a dataset of the most reliable dosage-insensitive (MRDIS) genes, we discarded the genes with a pLI value > 0.5 in either the dataset of Shihab et al. (2017) or the dataset of Lek et al. (2016) from these 9459 genes. In total, 5579 MRDIS genes were obtained, including 368 TF genes (Supplementary Table 2). Statistical analysis showed that TF genes were not significantly overrepresented or significantly underrepresented among the MRDIS genes (chi-square = 1.4, df = 1, BH-adjusted *p* = 0.569). These 368 MRDIS TF genes were distributed in 25 TF families (Table 2). Among the MRDIS TFs, members of the gene families C2H2-ZF and CENPB were significantly overrepresented, with a greater incidence than expected (BH-adjusted *p* < 0.05 for both cases). In the Homeodomain gene family, significantly fewer MRDIS genes (15) were observed than expected (47.9) (chi-square = 17.2, df = 1, BH-adjusted *p* = 0.001). No nuclear receptor genes were observed in the MRDIS gene dataset, but the expected value by chance was 11.3. Thus, nuclear receptor genes were significantly underrepresented in the MRDIS gene dataset (chi-square = 11.3, df=1, BH-adjusted *p* = 0.009). In all other TF gene families, the numbers of TF members were not significantly different from the expected values (BH-adjusted *p* > 0.05 for all cases). The small TF gene families were underrepresented in the MRDIS gene dataset regardless of whether they were defined by ≤ 5, ≤ 7, ≤ 9, or ≤ 11 members (BH-adjusted *p* < 0.05 for all cases, Table 2).

**Table 2.** TF genes in the most reliable dosage-insensitive (MRDIS) gene dataset.

| TF family | Total number in human genome | Observed number of MRDS TFs | Expected number in this dataset | BH-adjusted $p$ |
| --- | --- | --- | --- | --- |
| C2H2-ZF | 747 | 281 | 183 | $4 \times 10^{-4}$ |
| Homeodomain | 196 | 15 | 48 | 0.001 |
| Nuclear receptor | 46 | 0 | 11 | 0.009 |
| CENPB | 11 | 8 | 2.7 | 0.010 |
| All TFs | 1639 | 368 | 401 | 0.569 |
| Small-family TFs |  |  |  |  |
| $\leq 5$ members | 87 | 6 | 21 | 0.030 |
| $\leq 7$ members | 126 | 8 | 31 | 0.004 |
| $\leq 9$ members | 186 | 15 | 45 | 0.002 |
| $\leq 11$ members | 238 | 27 | 58 | 0.009 |

The chi-square test (expected value $> 5$ ) and Fisher's exact test (expected value $\leq 5$ ) were used to test the overrepresentation or underrepresentation of the TF genes in this dataset. The Benjamini-Hochberg (BH) procedure was used to compute the false discovery rate-adjusted $p$ values. The MRDIS genes are the genes with pLI values $< 0.5$ in either the dataset of Shihab et al. (2017) or the dataset of Lek et al. (2016) that were not considered as dosage-sensitive genes in any of the four datasets obtained by Shihab et al. (2017), Lek et al. (2016), Makino et al. (2013) and Rice and McLysaght (2017b).

Furthermore, we surveyed the abundance of KZFPs, which form a major subfamily of C2H2-ZF, in the above two datasets separately. Among the MRDS genes, there were no KZFP genes, although the expected value by chance was 14 (chi-square = 14.4, df = 1, *p* = 1.5 × 10^−4^). Furthermore, the KZFP genes were significantly overrepresented in the MRDIS gene dataset (observed/expected = 209/94, chi-square = 43.7, df = 1, *p* = 3.8 × 10^−11^).

### 3.2 MRDS TF Genes Are Enriched for a Variety of GO Terms, but MRDIS TF Genes Are Enriched for Only Three

GO enrichment analysis showed that the MRDS TFs were significantly enriched for 43 molecular function terms, 10 cellular component terms and 581 biological process terms (Figure 2 and Supplementary Table 3-5), consistent with the various roles of the TFs in regulating gene expression. In sharp contrast, the MRDIS TFs were significantly enriched for only three molecular function terms: 1) DNA-binding transcription repressor activity, RNA polymerase II-specific; 2) DNA-binding transcription activator activity, RNA polymerase II-specific; and 3) TF activity, RNA polymerase II proximal promoter sequence-specific DNA binding (Figure 2 and Supplementary Table 3-5). For each term, the matched TFs accounted for fewer than 10% of the MRDIS TFs; the majority of MRDIS genes did not significantly match any GO terms. These three GO terms for which MRDIS TFs were enriched were also the significant GO terms for the MRDS TFs (Figure 2).

**Figure 2.**
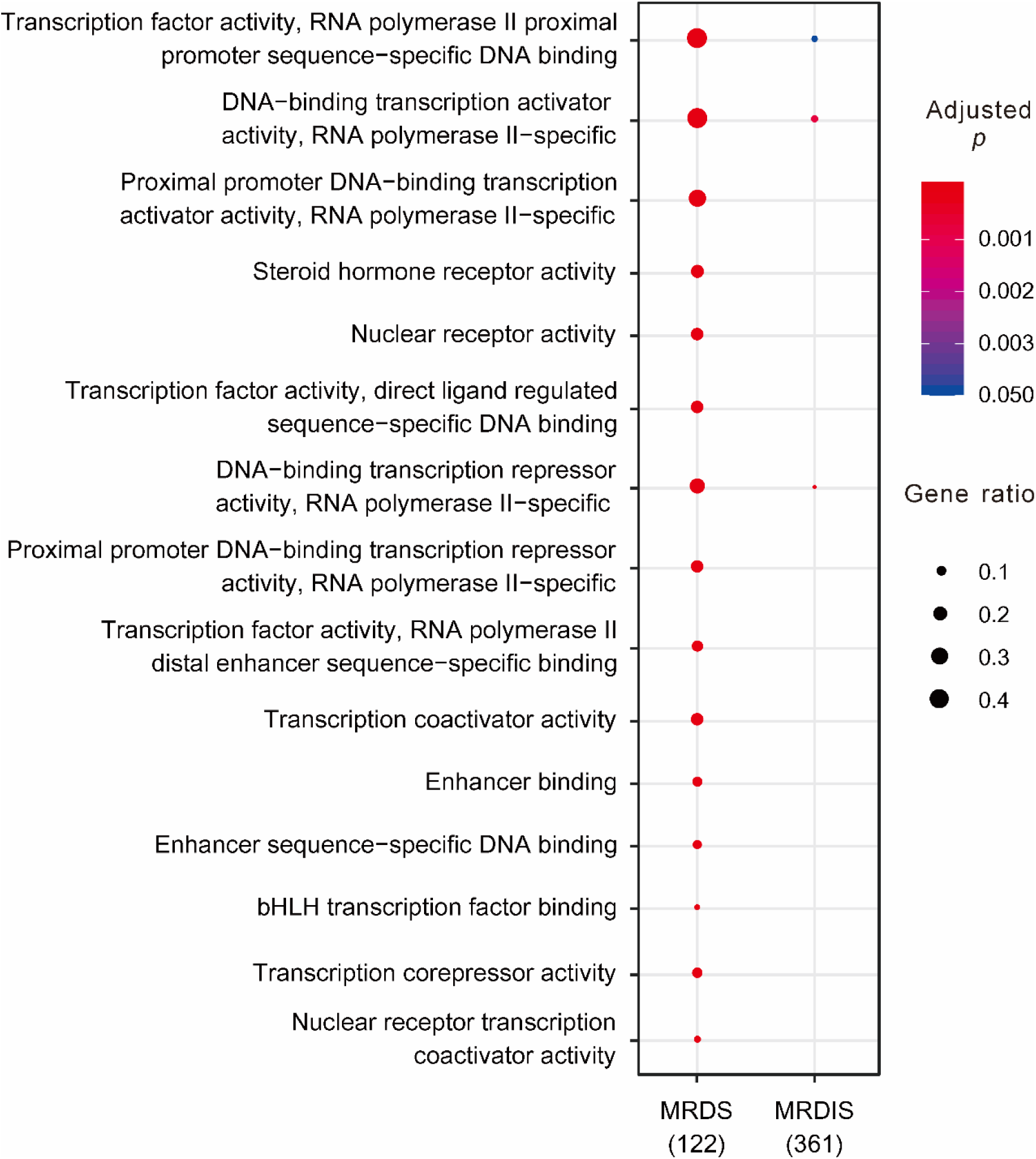
Clear differences in the Gene Ontology enrichment between MRDS TFs and MRDIS TFs. Due to space limitations, only the most significant terms of the MRDS TFs are displayed. The color of each circle represents the significance of the enrichment adjusted using the Benjamini-Hochberg (BH) procedure, and the size of each circle represents the percentage of genes associated with that term.

### 3.3 MRDS TFs Interact with More Proteins than MRDIS TFs

We downloaded human PPI data from the STRING database (Szklarczyk et al., 2018). The most recent version of STRING, 11.0, contains PPI data for 19566 human proteins, including 121 MRDS TFs and 356 MRDIS TFs. On average, each MRDS TF was found to interact with 851 proteins, while each MRDIS TF was found to interact with 263 proteins. Mann-Whitney *U* tests showed that the difference was statistically significant (*p* < 10^−6^). In addition, we observed that 25% of the proteins interacting with the MRDS TFs were TFs while 17% of the proteins interacting with the MRDIS TFs were TFs (Mann-Whitney *U* test, *p* < 10^−6^).

Next, we performed GO enrichment analysis of the proteins interacting with the MRDS TFs and MRDIS TFs. Generally, the results were consistent with the commonly believed functions of TFs in gene expression regulation (Supplementary Table 6-8). Both the proteins interacting with MRDS TFs and those interacting with MRDIS TFs were enriched for numerous terms indicating gene expression regulation, such as 0000982 (TF activity, RNA polymerase II proximal promoter sequence-specific DNA binding), 0007389 (pattern specification process), 0005667 (TF complex). In addition, there were some differences in the ranks of the enriched terms between the two groups of proteins. Among the top 20 enriched terms, the two groups shared 15, 11, and 12 common terms in the molecular function, biological process, and cellular component categories, respectively. However, it is difficult to draw conclusions on the functional differences between MRDS TFs and MRDIS TFs based on their interacting proteins.

The GO enrichment analyses of the MRDIS TFs and their interacting proteins indicated entirely different functional roles. One possible explanation is that the MRDIS TFs are more poorly annotated in the GO database than their interacting proteins. As the functions of most KZFPs are unknown (Imbeault et al., 2017), KZFPs are likely under-annotated in the GO database. From the dataset of Imbeault et al. (2017), we retrieved 399 KZFPs. None of these KZFPs was found in the MRDS TF dataset, while 209 KZFPs were found in the MRDIS TF dataset. GO enrichment analysis showed that the MRDIS KZFPs were not significantly enriched for any GO terms (BH-adjusted *p* > 0.05 for all the terms in the molecular function, biological process, and cellular component categories). In the STRING database (Szklarczyk et al., 2018), we found PPI records for all 208 MRDIS KZFPs. In total, the proteins interacting with the 208 MRDIS KZFPs were significantly enriched for 130 molecular function terms, 1299 biological process terms, and 139 cellular component terms (BH-adjusted *p* < 0.05 for cases, Supplementary Table 9-11). The potential functions of the MRDIS KZFPs in regulating gene expression were evident from the most enriched GO terms of their interacting proteins, such as 0005667 (TF complex), 0044798 (nuclear TF complex), 0000790 (nuclear chromatin), 0090575 (RNA polymerase II TF complex), 0000982 (TF activity, RNA polymerase II proximal promoter sequence-specific DNA binding), 0001228 (DNA-binding transcription activator activity, RNA polymerase II-specific), 0001077 (proximal promoter DNA-binding transcription activator activity, RNA polymerase II-specific), 0001227 (DNA-binding transcription repressor activity, RNA polymerase II-specific), 0001078 (proximal promoter DNA-binding transcription repressor activity, RNA polymerase II-specific), 0003713 (transcription coactivator activity), 0003714 (transcription corepressor activity), 0001158 (enhancer sequence-specific DNA binding), 0035326 (enhancer binding), and 0070491 (repressing TF binding). These results indicate that some MRDIS TFs are under-annotated in the GO database. Furthermore, the Mann-Whitney *U* test showed that MRDIS KZFPs interact with significantly fewer proteins than other MRDIS TFs (average values: 204 *vs.* 346, *p* < 10^−6^), indicating that the under-annotated MRDIS TFs have lower functional complexity than the well-annotated ones.

### 3.4 Differences in the DBDs of MRDS TFs and MRDIS TFs

Protein domains are the significant tertiary structures of a protein that are generally believed to have more functional implications than the rest of the protein chain. A TF gene with two or more functions is more likely to be sensitive to dosage changes than one with fewer functions because a dosage constraint on any function will cause the gene to be dosage-sensitive. For this reason, we compared the number of DBDs between MRDS TFs and MRDIS TFs. The majority of human TFs had only one DBD in each protein. There were only 43 two-DBD TFs, including seven MRDS TFs and two MRDIS TFs. Although a significantly percentage of MRDS TFs than MRDIS TFs were two-domain proteins (chi-square = 4.30, df = 1, *p* = 0.038), the difference in dosage sensitivity between MRDS TFs and MRDIS TFs could not be attributed to the number of domains.

To test whether the dosage sensitivity of human TFs is achieved through dual functions of both activator and repressor properties, we searched the keywords “transcription factor,” “activator domain,” “repressor domain,” “activation domain,” and “repression domain” in the UniProtKB database (The UniProt Consortium, 2019). Among the 122 MRDS TFs, 21 TFs were found to have activation domains, while eight TFs were found to have repressor domains. Three MRDS TFs (*HIF1A, SP1*, and *TP63*) have both activation domains and repressor domains. Among the 368 MRDIS TFs, three have activation domains, and five have repressor domains. Two MRDIS TFs (*ZBTB32* and *YY2*) have both activation domains and repressor domains. According to Imbeault et al. (2017), the KRAB domains of KZFPs can repress transposable elements by recruiting transcriptional regulators such as TRIM28. Therefore, KRAB domains were also regarded as repressor domains in the present study. From Imbeault et al. (2017), we retrieved 399 KRAB-containing proteins, 209 of which were MRDIS TFs. None of these 209 TFs was found to have activation domains. However, MRDS TFs were found to be more likely to work as activators than MRDIS TFs (17.2% *vs.* 0.82%, chi-square = 14.9, *p* = 10^−4^), whereas MRDIS TFs are more likely to work as repressors (6.56% *vs.* 58.2%, chi-square = 41.1, *p* = 1.4 × 10^−10^ if regarding KRAB as a repressor domain; the difference was not significant if we did not regard KRAB as a repressor domain).

We also compared the sizes of DBDs between MRDS TFs and MRDIS TFs. Among the 77 MRDS TFs whose domains have been annotated, the average domain size was 118 amino acid residues. In contrast, the average domain size of the 279 MRDIS TFs was only 52 amino acid residues. Mann-Whitney *U* tests showed that the difference was highly significant (*p* = 2.1 × 10^−6^). Furthermore, we compared the sizes of the activation domains and the repressor domains. Regardless of whether the comparisons were performed within MRDS TFs, within MRDIS TFs, or within a combined dataset of MRDS TFs and MRDIS TFs, we did not detect any significant difference in either activation domain size or repressor domain size (Mann-Whitney *U* test, *p* > 0.05 for all cases).

### 3.5 Quaternary Structures of MRDS TFs and MRDIS TFs

We surveyed the quaternary structures of TFs using the data of Lambert et al. (2018) and found three types of proteins: low-specificity DNA-binding proteins, monomers or homomultimers, and obligate heteromers. The vast majority of the TFs were monomers or homomultimers. In total, 96.7% of MRDS TFs were monomers or homomultimers, and 97.6% of MRDIS TFs were monomers or homomultimers. Therefore, the dosage sensitivity of TFs is not related to the type of quaternary structure.

### 3.6 Differences in the DNA-binding Sites of MRDS TFs and MRDIS TFs

From Lambert et al. (2018), we obtained the DNA-binding sites of 102 MRDS TFs and 235 MRDIS TFs. On average, the DNA-binding sites of the MRDS TFs had 12 nucleotides, whereas those of the MRDIS TFs had 13 nucleotides. Although the difference seems slight, it was statistically significant (Mann-Whitney *U* test, *p* = 7.4 × 10^−5^, Table 3). Furthermore, we compared the nucleotide compositions of the DNA-binding sites. The DNA-binding sites of both MRDS TFs and MRDIS TFs had greater numbers of A than any of the other three nucleotides (Mann-Whitney *U* test, *p* < 10^−6^ for all comparisons). However, the DNA-binding sites of MRDS TFs had significantly fewer A residues than those of MRDIS TFs (Mann-Whitney *U* test, *p* < 10^−6^). Furthermore, the DNA-binding sites of MRDS TFs had more G residues than those of MRDIS TFs (Mann-Whitney *U* test, *p* = 0.048). No significant differences were observed in C and T.

**Table 3.**
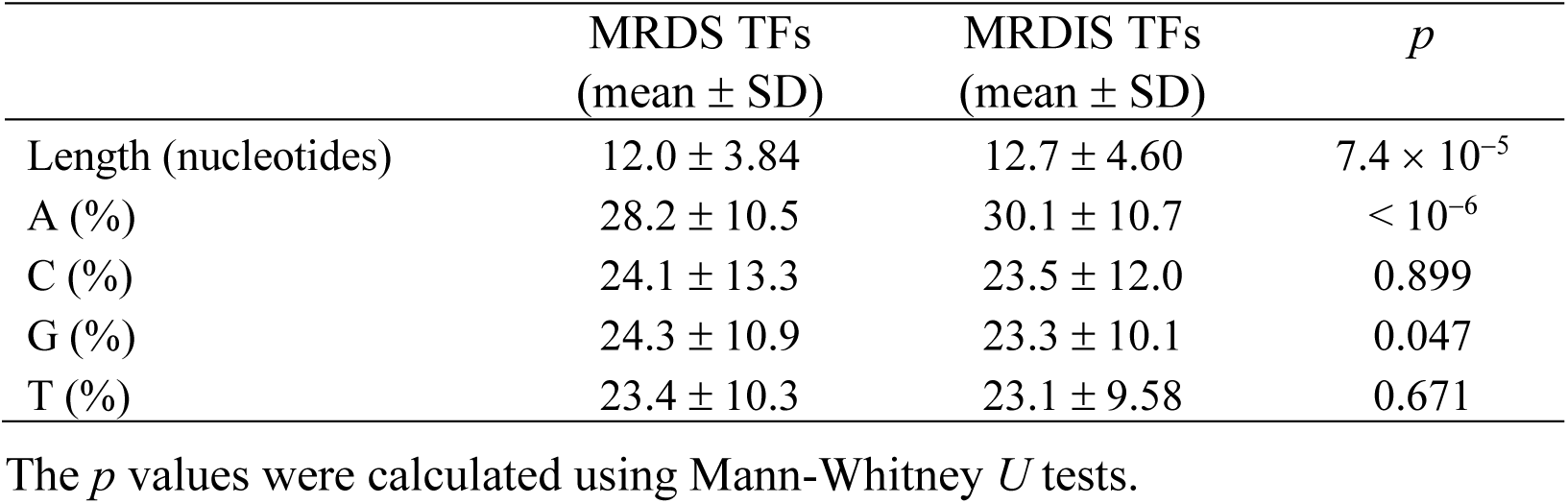
Differences in the DNA-binding sites of the MRDS TFs and MRDIS TFs.

### 3.7 Expression Patterns of MRDS TFs and MRDIS TFs

One previous study has shown that dosage-sensitive genes are generally expressed at high levels (Gout et al., 2010; Rice and McLysaght, 2017a). From the Human Protein Atlas (Uhlén et al., 2015), we retrieved the mRNA expression levels of 119 MRDS TFs and 348 MRDIS TFs in 37 cell samples. After averaging the mRNA expression levels of each TF among these 37 samples, we compared the expression levels between the 119 MRDS TFs and 348 MRDIS TFs. Consistent with previous studies, we found that the mRNA expression levels of MRDS TFs (6.36 ± 0.679) were significantly higher than those of MRDIS TFs (6.36 ± 0.679 *vs.* 6.26 ± 1.767; Mann-Whitney *U* test, *p* = 0.001).

Although mRNA abundance is significantly correlated with protein abundance, it is far from a perfect parameter to represent or predict protein expression levels (Greenbaum et al., 2003; Gry et al., 2009; Maier et al., 2009). Therefore, we re-examined the relationship between dosage sensitivity and gene expression level using protein abundance data. In the Human Protein Atlas database, the expression level of a protein, if it has ever been studied, is defined as high, medium, low, or not detected in each studied cell sample. To ensure that expression data existed for all the studied genes in the studied cell samples, we selected 31 cell samples in which 684 MRDS genes (including 92 MRDS TF genes) and 2736 MRDIS genes (including 150 MRDIS TF genes) had ever been studied. For each gene, we surveyed its expression level in the 31 cell samples. For example, the expression level of gene *ENSG00000095951*, a member of the C2H2-ZF gene family, was high in nine samples, medium in 12 samples, low in eight samples, and not detected in two samples. We first compared the presence of the MRDS genes and MRDIS genes in each of the four categories. On average, each MRDS gene had high expression levels in 4.5 samples, while each MRDIS had high expression levels in 3.0 samples. The Mann-Whitney *U* test showed that the difference was statistically significant (*p* < 10^−6^; Table 4). Similarly, at the medium and low levels, MRDS genes were detected in significantly more samples than MRDIS genes (Mann-Whitney *U* test, *p* < 10^−6^ for both cases; Table 4). In the “not detected” category, MRDS genes were assigned to significantly fewer samples than MRDIS genes (Mann-Whitney *U* test, *p* < 10^−6^; Table 4). However, no significant differences were detected between MRDS TF genes and MRDIS TF genes in any of the four categories (Mann-Whitney *U* test, *p* > 0.05 for all cases). Thus, we found that dosage-sensitive genes in general have higher expression levels than dosage-insensitive genes, confirming the findings of a previous study. However, our results suggest that dosage-sensitive TF genes do not have higher expression levels than dosage-insensitive TF genes.

**Table 4.** Differences in the protein expression patterns of the MRDS TFs and MRDIS TFs.

|  | MRDS genes | MRDIS genes | *p* | MRDS TFs | MRDIS TFs | *p* |
| --- | --- | --- | --- | --- | --- | --- |
| Number of genes | 684 | 2736 |  | 92 | 150 |  |
| Number of cell samples in which the proteins are |  |  |  |  |  |  |
| at high levels | 4.5 ± 6.6 | 3.1 ± 4.8 | < 10<sup>-6</sup> | 4.6 ± 6.6 | 3.8 ± 5.4 | 0.520 |
| at medium levels | 9.2 ± 6.6 | 7.6 ± 6.7 | < 10<sup>-6</sup> | 9.3 ± 7.2 | 9.8 ± 6.2 | 0.416 |
| at low levels | 5.9 ± 4.3 | 4.8 ± 4.2 | < 10<sup>-6</sup> | 5.4 ± 4.3 | 6.2 ± 4.4 | 0.116 |
| not detected | 11.3 ± 9.9 | 15.5 ± 11.1 | < 10<sup>-6</sup> | 11.8 ± 10.8 | 11.1 ± 9.5 | 0.960 |

In total, 31 cell samples were studied for each gene. The MRDS genes and the MRDIS genes are defined in Tables 1 and 2 as well as in the main text. The *p* values were calculated using Mann-Whitney U tests.

### 3.8 Protein Size Differences between MRDS TFs and MRDIS TFs

Rice and McLysaght (2017a) found that ohnologs have much longer coding sequences than genes duplicated by small-scale duplication. We re-examined this phenomenon using our datasets. In version GRCh37.p13 of the human genome, we obtained the coding sequences (CDSs) of 5569 (among the 5579) MRDIS genes. The CDSs of the 853 MRDS genes were significantly longer than those of the MRDIS genes (Mann-Whitney *U* test, *p* < 10^−6^), with a difference in average values of more than 2-fold (2640 *vs*. 1310, Table 5). The MRDS TF genes and the MRDIS TF genes also differed significantly in CDS length (Mann-Whitney *U* test, *p* = 10^−6^), but there was a smaller difference in their average values (2189 *vs*. 1577, Table 5). Furthermore, we noticed that TF genes had significantly longer CDSs than other protein-coding genes (Mann-Whitney U test, *p* < 10^−6^, Table 5).

**Table 5.** Comparison of coding sequence lengths.

|  | Number of genes | Mean ± SD (bp) | *p* |
| --- | --- | --- | --- |
| MRDS genes | 853 | 2640 ± 1783 | < 10<sup>-6</sup> |
| MRDIS genes | 5569 | 1310 ± 1956 |  |
| MRDS TFs | 122 | 2189 ± 1444 | 10<sup>-6</sup> |
| MRDIS TFs | 368 | 1577 ± 698 |  |
| TFs | 1608 | 1777 ± 1205 | < 10<sup>-6</sup> |
| Other genes | 19634 | 1710 ± 1826 |  |

The MRDS genes and the MRDIS genes are defined in Tables 1 and 2 as well as in the main text. The *p* values were calculated using Mann-Whitney *U* tests.

### 3.9 MRDS TF Genes Experience Stronger Selective Pressures than MRDIS Genes

Schuster-Böckler et al. (2010) showed that dosage-sensitive genes are under strong selective force. In this study, we used the *d*_N_/*d*_S_ value for each human gene relative to its mouse ortholog as a measure of the selective pressure of each gene. Valid *d*_N_/*d*_S_ values were obtained for 4188 MRDIS genes and all 853 MRDS genes. Mann-Whitney *U* tests showed that the MRDS genes experienced significantly stronger selective pressure than the MRDIS genes (*p* < 10^−6^, Table 6). The MRDS TF genes and MRDIS TF genes exhibited the same patterns (Mann-Whitney *U* test, *p* < 10^−6^, Table 6). Compared with other genes in the human genome, TF genes are under stronger selective forces (Mann-Whitney *U* test, *p* < 10^−6^, Table 6).

**Table 6.** Comparison of selective pressures.

|  | Number of genes | *d<sub>N</sub>/d<sub>S</sub>* ± SD | *p* |
| --- | --- | --- | --- |
| MRDS genes | 853 | 0.062 ± 0.057 | < 10<sup>-6</sup> |
| MRDIS genes | 4188 | 0.257 ± 1.536 |  |
| MRDS TFs | 122 | 0.066 ± 0.054 | < 10<sup>-6</sup> |
| MRDIS Fs | 226 | 0.256 ± 0.217 |  |
| TFs | 1374 | 0.134 ± 0.141 | < 10<sup>-6</sup> |
| Other genes | 16804 | 0.161 ± 0.776 |  |

The MRDS genes and the MRDIS genes are defined in Tables 1 and 2 as well as in the main text. The *p* values were calculated using Mann-Whitney U tests.

### 3.10 HLOF-Tolerant TF Genes Are Not Rarer than Other Human Genes

It has been determined experientially that complete homozygous loss of TF genes rarely occurs in humans (Engelkamp and van Heyningen, 1996). Here, we used recent datasets of human HLOF-tolerant genes to examine whether TF genes are sparser than other genes in the human genome. A series of large-scale sequencing studies have been carried out on healthy human adults to identify HLOF-tolerant genes (Lim et al., 2014; Sulem et al., 2015; Lek et al., 2016; Narasimhan et al., 2016; Saleheen et al., 2017; Bartha et al., 2018). We retrieved the data from the two most extensive studies (Lek et al., 2016; Saleheen et al., 2017). In addition to the abovementioned pLI metric, Lek et al. (2016) designed another metric, pNull. With a cutoff of a pNull > 0.9, 1226 genes were identified as extremely loss of function-tolerant from their dataset. In version GRCh37.p13 of the human genome, 22810 protein-coding genes have been annotated; thus, 5.4% of human genes are HLOF-tolerant. Among the 1639 TF genes, we found 103 genes (6.3%) in this HLOF-tolerant gene dataset. The TF genes were not significantly over- or underrepresented among the HLOF-tolerant genes (BH-adjusted *p* value = 1). However, we found that one TF gene family, C2H2-ZF, was significantly enriched in HLOF-tolerant genes (observed value = 91, expected value =40, BH-adjusted *p* value = 0.0006). Furthermore, we found that KZFPs, a subfamily of C2H2-ZF, had an even higher observed/expected ratio (81/21, chi-square = 34.8, df = 1, *p* = 3.6 × 10^−9^). In a more recent study, Saleheen et al. (2017) sequenced the protein-coding regions of 10503 adult participants with a high rate of consanguinity and identified 1317 distinct genes for which loss-of-function mutations in both copies were tolerated, including 85 TF genes. Statistical analysis did not show significant over- or underrepresentation of TF genes among the HLOF-tolerant genes (expected value = 104, chi-square = 1.9, BH-adjusted *p* value = 1). The same result was observed when the enrichment of each TF gene family (except for KZFPs, a subfamily of C2H2-ZF) was analyzed separately (BH-adjusted *p* value > 0.05 for all cases). The observed number of KZFPs in this HLOF-tolerant gene dataset was more than twice the value expected by chance (55 vs. 22, chi-square = 14.0, df = 1, p = 1.9 × 10^−4^).

## 4 DISCUSSION

TFs are a group of proteins with unique functional roles to ensure the temporally and spatially accurate expression of the genetic information encoded in the genome. Intuitively, changes in their gene copy numbers should be hazardous because changes in the concentrations of TFs in cells will disturb the normal expression patterns of the TF target genes. Consistent with this idea, complete homozygous loss of TF genes is rarely observed in humans (Engelkamp and van Heyningen, 1996). Using the two most extensive datasets of human HLOF-tolerant genes (Lek et al., 2016; Saleheen et al., 2017), we found that TF genes are not significantly underrepresented among HLOF-tolerant genes. At least with regard to their percentages, TF genes are not more indispensable than other genes in the human genome. However, a specific TF family, C2H2-ZF, seems to be more dispensable than other genes in the human genome. In particular, the genes of the large KZFP subfamily, which accounts for 51% of the C2H2-ZF gene family, are significantly more dispensable than other genes in the human genome. Imbeault et al. (2017) showed that the majority of KZFPs bind transposable elements and repress these transposable elements rather than regulating the expression of other protein-coding genes. With the rapid accumulation and decay of transposable elements during evolution (Blass et al., 2012; Wallau et al., 2014), the KZFP subfamily has also likely experienced dynamic gain of new members and loss of obsolete members.

Compared with complete homozygous losses, gain of a new copy or heterozygous loss of one gene copy in a diploid genome is expected to have a gentler effect. Seidman and Seidman (2002) surveyed the haploinsufficiency of human TFs using the data available at that time. Deleterious phenotypic effects resulting from heterozygous losses were confirmed in only 27 of the 491 TF genes. From four recent datasets on human dosage-sensitive genes, including datasets obtained from both heterozygous loss studies and gene duplication studies (Makino et al., 2013; Lek et al., 2016; Rice and McLysaght, 2017b; Shihab et al., 2017), we defined a dataset of the most reliable dosage-sensitive genes, the MRDS genes, by selecting the dosage-sensitive genes shared by all four datasets. As a control, we also defined a dataset of the most reliable dosage-insensitive genes, the MRDIS genes, by selecting the genes that were not considered dosage-sensitive in any of the four datasets. Upon surveying the abundances of the TF genes in these two datasets, we observed some commonalities (Table 1-2).

First, TF genes were more likely to be dosage-sensitive than other genes in the human genome. The significant enrichment of TF genes in the dosage-insensitive gene datasets was expected given the roles of TFs in ensuring accurate expression of their target genes.

Second, the nuclear receptor is a very unique TF gene family with regard to dosage sensitivity. Members of this family were significantly overrepresented in the MRDS gene dataset and absent from the MRDIS gene dataset. Nuclear receptors can activate target genes after binding nonpolar ligands such as estrogen, progesterone, retinoic acid, oxysterols, and thyroid hormone. In contrast to other TFs, nuclear receptors respond directly to extracellular changes by binding ligands that are diffusible across the plasma membrane (Sladek, 2011). A distinguishing characteristic of steroid hormones is the dose-response curve manifested in their regulation of gene expression (Szapary et al., 1999; Simons, 2006). Fixed dosages of the nuclear receptors for steroid hormones might be the premise of the dose-response curve.

Third, TF gene families with fewer numbers were more likely to be enriched among the dosage-sensitive genes. This finding was supported by observations of overrepresentation of small-family TF genes in the MRDS gene dataset and underrepresentation of them in the MRDIS gene dataset. We did not observe any common characteristics among these small-family TFs. Instead, we suspect that the dosage sensitivity of these TF genes might act as a selective force against the expansion of these families and maintain the small number of members during evolution.

Fourth, many TF genes were dosage-insensitive, and the KZFP genes seemed to be the most dosage-insensitive. Considering the roles of TF genes in the regulation of gene expression, the existence of a large number of dosage-insensitive TF genes was unexpected. One possible explanation is that some TFs do not directly regulate the expression levels of other genes but simply contribute, more or less, to the maintenance of a nuclear environment compatible with the expression of required genes. Nonfunctional interactions within the accessible portions of the genome have been observed for many TFs (Fisher et al., 2012; Slattery et al., 2014). Some TFs, such as KZFPs, have been shown to be involved in the repression of transposable elements (Thomas and Schneider, 2011; Imbeault et al., 2017; Yang et al., 2017). As there is no purifying selection against the accumulation of degenerative mutations in repressed transposable elements, most transposable elements quickly lose their transposition potential during evolution. Dosage changes in TFs that target transposable elements with low or no transposition potential should not have severe effects. In addition, if large proportions of TF binding events are nonfunctional, the effects of dosage changes in TF genes on the expression of target genes should be buffered. Nonfunctional binding (Todeschini et al., 2014) might therefore underlie the mechanism of dosage insensitivity of some TF genes.

Then, we characterized the dosage-sensitive TFs by comparing the MRDS TFs and the MRDIS TFs.

GO enrichment analysis showed that MRDS TFs were significantly enriched for 634 GO terms, whereas MRDIS TFs were significantly enriched for only three molecular function terms (Supplementary Table 3-5), indicating that MRDS TFs have a much greater variety of functions than MRDIS TFs. Consistent with this result, we found that MRDS TFs interact with many more proteins than MRDIS TFs. GO enrichment analysis of the proteins interacting with MRDS TFs and MRDIS TFs, however, showed that both groups of proteins were enriched for numerous terms indicating gene expression regulation. The KZFP family is a unique TF family; no member of this family fell in the MRDS category. The MRDIS KZFPs were not significantly enriched for any GO terms, whereas the proteins found to interact with the MRDIS KZFPs were significantly enriched for thousands of GO terms (Supplementary Table 9-11). The most enriched GO terms of the proteins interacting with the MRDIS KZFPs provide evidence that the MRDIS KZFPs are actively involved in regulating gene expression. In addition, we found that MRDIS KZFPs interact with fewer proteins than other MRDIS TFs. It might be concluded that KZFPs and perhaps some other MRDIS TFs have lower functional complexity than MRDS TFs; however, these TFs are under-annotated in the GO database compared with MRDS TFs.

In plants, the TF WUS provides an insightful example of the mechanism of dosage sensitivity of TFs (Hofhuis and Heidstra, 2018). WUS can bind different cofactors and activates its target gene CLV3 when its protein is expressed at low levels, while it represses the target gene when its protein is at high levels. For the WUS gene, both an increase and a decrease in its dosage would disturb its function. The dosage of the WUS gene has been fixed by its dual functions as an activator and a repressor. We speculated that these dual functions could be achieved by either a protein complex containing both activator subunits and repressor subunits or a protein containing both repressor domains and activation domains. Our survey of the quaternary structures of TFs showed that the vast majority of both MRDS TFs and MRDIS TFs are monomeric or homomultimeric rather than heteromeric. Furthermore, we found that the majority of human TFs have a single DBD. For a few MRDS TFs and a few MRDIS TFs, both activation domains and repressor domains have been annotated in the UniProtKB database (The UniProt Consortium, 2019). However, the activation or repression functions of most TFs have not yet been annotated in the database; therefore, more data is needed before a solid conclusion can be made.

The DNA-binding sites of MRDS TFs were shorter and contained fewer A and slightly more G residues than those of the MRDIS TFs.

Previous analyses have characterized dosage-sensitive genes as being highly expressed, encoding large proteins, and being under strong selective force (Gout et al., 2010; Schuster-Böckler et al., 2010; Rice and McLysaght, 2017a). In this study, we confirmed most of these conclusions by comparing MRDS genes and MRDIS genes as well as MRDS TF genes and MRDIS TF genes; however, we found that MRDS TFs do not have significantly higher protein expression levels than MRDIS TFs. Dosage sensitivity could not be explained by expression burden, at least among the TF genes. The small *d*_N_/*d*_S_ values and the consequently high selective pressures experienced by the dosage-sensitive genes indicate that functional constraints might be tightly associated with dosage constraints.

Using the dosage-sensitive gene and dosage-insensitive gene datasets we compiled based on four recent datasets of dosage-sensitive genes, we found that human TFs were significantly enriched among the dosage-sensitive genes. In addition, many TFs were found to be dosage-insensitive. The most dosage-sensitive TF gene family was the nuclear receptor family, while the most dosage-insensitive TF gene family was the KZFP subfamily. Further characterization of these genes revealed both intrinsic differences between the dosage-sensitive genes and the dosage-insensitive genes and the relatively limited knowledge regarding some dosage-insensitive genes, such as KZFPs.

## 6 Author Contributions

DKN conceived the study and wrote the manuscript. ZN retrieved the data from online databases and performed all the statistical analyses. XYZ and SA each repeated some of the analyses. All authors read and approved the final manuscript.

## 8 Acknowledgments

We appreciate the helpful comments from the handling editor and the anonymous referees, and the kind helps from Er-Li Pang and the High-Performance Computing Center of Hebei University. This manuscript has been released as a preprint at BioRxiv (Ni et al., 2019).

